# Vitamin B12 supports skeletal muscle oxidative phosphorylation capacity in male mice

**DOI:** 10.1101/2025.05.19.654973

**Authors:** Luisa F. Castillo, Katarina E. Heyden, Abigail R. Williamson, Wenxia Ma, Olga V. Malysheva, Nathaniel M. Vacanti, Anna E. Thalacker-Mercer, Martha S. Field

**Author notes:** **Corresponding author:** Martha S. Field. These authors contributed equally to this work (Luisa F. Castillo, Katarina E. Heyden).

## Abstract

**Objectives:** Vitamin B12 plays a vital role in folate-mediated one-carbon metabolism (FOCM), a series of one-carbon transfer reactions that generate nucleotides (thymidylate (dTMP) and purines) and methionine. Inadequate levels of B12 impair FOCM, depressing *de novo* thymidylate (dTMP) synthesis, which in turn leads to uracil accumulation in DNA. This phenomenon has been well documented in nuclear DNA. Our previous work in liver tissue has shown that mitochondrial DNA (mtDNA) is more sensitive to FOCM impairments in that mtDNA exhibits elevated uracil levels before uracil concentrations in nuclear DNA change. However, the functional consequences of uracil accumulation in mtDNA are largely unknown. The purpose of this study was to determine how a functional B12 deficiency (induced by reduced levels of the B12-dependent enzyme methionine synthase (MTR)) and dietary B12 deficiency affects mtDNA integrity and mitochondrial function in energetic and mitochondria-rich tissues such as skeletal muscle.

**Methods:** Male *Mtr^+/+^* and *Mtr^+/−^* mice were weaned to either an AIN93G-based control (C) diet containing 25 µg/kg vitamin B12 or a B12-deficient (-B12) diet containing 0 µg/kg vitamin B12 to explore the effects of functional (*Mtr^+/−^*) and dietary B12 deficiency on muscle weight, uracil content in mtDNA, mtDNA content, and oxidative phosphorylation complex capacity in skeletal muscle. Aged (20-22mo) male C57BL6/N mice were acclimated to an AIN93G control diet four weeks, then received either weekly injections of saline (vehicle control [30 uL 0.9% NaCl]) or B12 (0.65mg per 30uL 0.9% NaCl) in each of two hindleg muscles [1.25 mg B12 total]) for 8 weeks.

**Results:** The tibialis anterior (TA) muscle from *Mtr^+/−^*mice exhibited lowered maximal respiratory capacity of complex I, II, and IV of the electron transport chain than did TA from *Mtr^+/+^* mice. Exposure to the -B12 diet lowered maximal capacity of complex I in red, mitochondrially rich muscle (soleus and mitochondria-rich portions of quadriceps and gastrocnemius) (p=0.02). Levels of uracil accumulation in mtDNA in red muscle and gastrocnemius were elevated ∼10 fold with exposure to -B12 diet (p=0.04 and p<0.001, respectively). In aged mice gastrocnemius complex IV activity increased with intramuscular B12 supplementation (p=0.04)

**Conclusions:** Exposure to a B12-deficient diet led to uracil accumulation in mtDNA and impaired maximal oxidative capacity in two different types of skeletal muscle. B12 supplementation improved complex IV maximal capacity in gastrocnemius from aged mice.

## Introduction

Vitamin B12 (B12) deficiency disproportionately affects subpopulations including older adults and vegans/vegetarians (1). This deficiency commonly manifests through hematological and neurological symptoms due to the role of B12 in supporting DNA synthesis. B12 is an essential cofactor required for folate-mediated one-carbon metabolism (FOCM), a metabolic pathway which provides one-carbon groups for *de novo* biosynthesis of nucleotides (thymidylate (dTMP) and purines) and methyl donor generation (1) . B12 is required as a cofactor by only two enzymes in the body: methyl malonyl CoA mutase (MCM), which supports metabolism of amino acids and fatty acids through the tricarboxylic acid cycle and resides in the mitochondria, and methionine synthase (MTR), which is part of FOCM and localizes to the cytosol (1).

MTR requires B12 and folate to catalyze regeneration of methionine from homocysteine. In this two-step process, 5-methyl-tetrahydrofolate (5-methyl-THF) donates a methyl group to cobalamin (a form of B12), releasing THF and generating methylcobalamin (2). Methylcobalamin then provides the methyl group to homocysteine for methionine synthesis (2). The conversion of 5,10-methylene-THF to 5-methyl-THF by methylenetetrahydrofolate reductase (MTHFR) is irreversible, and the only enzyme capable of metabolizing 5-methyl-THF to release THF is MTR (**Figure 1**) (3). In instances of B12 deficiency or reduced *Mtr* expression, folate becomes increasingly trapped as 5-methyl-THF, which decreases the availability of other folate cofactors (i.e., THF and 5,10-methylene-THF) needed for nucleotide synthesis including synthesis of thymidylate (dTMP, of the “T” base in DNA) (Figure. 1).

**Figure 1.**
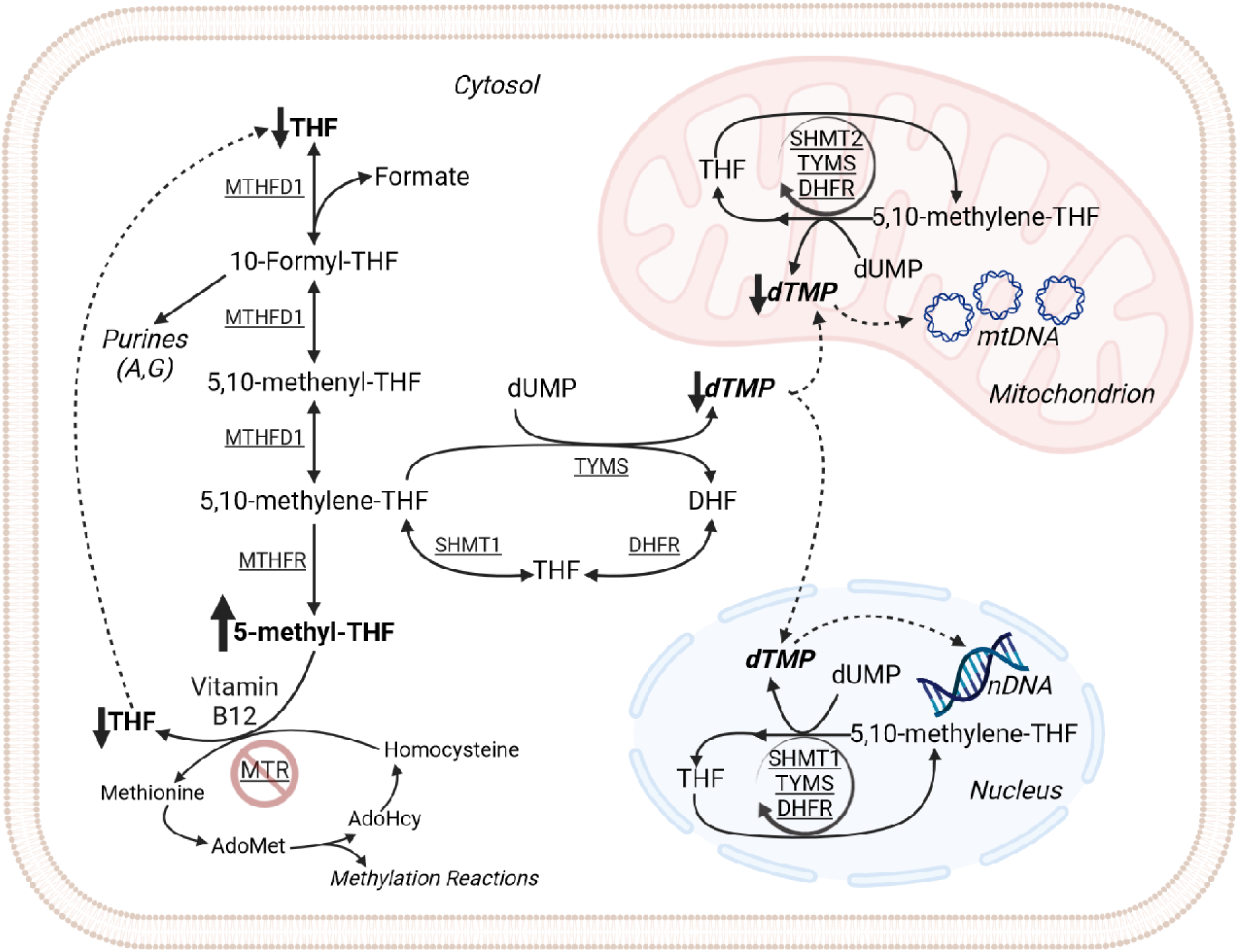
The effects of decreased *Mtr* expression on folate-mediated one-carbon metabolism. Vitamin B12 deficiency, as modeled by partial knockout of the *Mtr* gene in mice causes elevation of cellular folate as 5-methyl-THF and matched reduction of folate as THF. The conversion of 5,10-methylene-THF to 5-methyl-THF is irreversible, and 5-methyl-THF can only be metabolized to THF by the MTR enzyme using B12 as a cofactor. AdoHcy, S-adenosylhomocysteine; AdoMet, S-adenosylmethionine; DHFR, dihydrofolate reductase; dTMP, deoxythymidine monophosphate; dUMP, deoxyuridine monophosphate; FOCM, folate-mediated one-carbon metabolism; MTHFD1, methylenetetrahydrofolate dehydrogenase 1; MTHFR, methylenetetrahydrofolate reductase; MTR, methionine synthase; mtDNA, mitochondrial DNA; nDNA, nuclear DNA; SHMT, serine hydroxymethyltransferase; THF, tetrahydrofolate; TYMS, thymidylate synthase.

When dTMP levels are low, DNA polymerases incorrectly incorporate a uracil (or “U”) base into DNA in place of “T” (4). Uracil misincorporation initiates base-excision repair mechanisms, which in the continued absence of dTMP, lead to DNA double-strand breaks, stalled replication fork progression, and DNA instability (4). Folate-dependent dTMP synthesis occurs in multiple cellular compartments (cytosol, nucleus, and mitochondria) (5,6). The impact of FOCM impairments on DNA stability has been almost exclusively studied in the context of nuclear DNA (nDNA). However, there is evidence that impaired FOCM, either by dietary or genetic means, leads to decreased dTMP synthesis and uracil misincorporation in mitochondrial DNA (mtDNA) as well as nDNA (5,7,8).

mtDNA, though relatively small (16.5 kb), codes for several proteins that comprise the electron transport chain complexes in the mitochondria. Expression of these proteins and their complexes is crucial for cellular energy production via oxidative phosphorylation (OXPHOS) (9). mtDNA replication is one component of mitochondrial biogenesis, and maintenance of mtDNA is closely tied to mitochondrial function (9). Nucleotide synthesis, specifically *de novo* dTMP synthesis, has been shown to occur in the mitochondria and to be essential for maintaining mtDNA integrity (6). We have previously shown in mouse liver that mtDNA is more sensitive to FOCM impairments and exhibits elevated uracil levels before nDNA (7,8). However, the functional consequences of uracil accumulation in mtDNA are largely unknown. To address this gap, the present study aimed to investigate mitochondrial DNA integrity and mitochondrial function under B12-deficient conditions by genetically disrupting the B12-dependent *Mtr* enzyme in mice and inducing dietary B12 deficiency. Given that tissues with high energy demands and abundant mitochondria are particularly susceptible to mitochondrial dysfunction, we focused our analysis on skeletal muscle, hypothesizing that it would be especially vulnerable to the effects of B12 deficiency. Furthermore, we sought to examine whether dietary B12 supplementation could mitigate age-associated changes mitochondrial function.

## Methods

### Mouse housing and husbandry for *Mtr^+/−^* study

The *Mtr^+/−^* mouse model, generated as previously described (10), was backcrossed for over 10 generations and crossed to C57BL/6J. Male *Mtr^+/+^* and *Mtr^+/−^*offspring were randomly weaned to one of two defined diets at three weeks of age. The diets consisted of defined AIN93G-based control (C) diet containing 25 µg/kg vitamin B12 (#117883GI; Dyets, Inc., Bethlehem, PA), or an AIN93G-based vitamin B12 deficient (-B12) diet containing 0 µg/kg vitamin B12 (#117884GI; Dyets, Inc., Bethlehem, PA). Both diets included 1% succinyl sulfathiazole antibiotic to facilitate B12 deficiency by diminishing the contribution of B12 from gut microbes. The body weights of the mice were recorded at 0-, 2-, and 4-weeks post-weaning. Mice were maintained on the assigned diets for 7 weeks and sacrificed by cervical dislocation following CO_2_ euthanasia. Whole blood was collected via cardiac puncture into Heparin-coated tubes. Plasma and red blood cells were separated by centrifugation at 2500 x g and immediately flash frozen in liquid nitrogen. Mitochondria were isolated from red, mitochondria rich skeletal muscle tissue (cumulation/sum of the soleus as well as the red portions from the quadriceps and gastrocnemius) as previously described (11). Quadricep, tibialis anterior, gastrocnemius, and soleus wet wights were normalized to tibia length (mm). All other tissues were harvested, rinsed in ice-cold 1X phosphate buffered saline (PBS, Corning), and immediately flash-frozen in liquid nitrogen. All samples were then stored at −80°C for further analysis.

### Mouse housing and husbandry for aged C57BL/6N mice

Aged (20-22mo) C57BL6/N male mice were received from Charles River Laboratories. Before starting treatment, mice were acclimated to the defined AIN93G control diet for four weeks. Mice were then assigned randomly to either control- or B12-supplemented treatment. Control mice received saline injections (30 mL 0.9% NaCl) and a supplemented mice received B12 injections (0.65mg per 30mL 0.9% NaCl [saline] in each of two hindleg muscles [1.25 µg B12 total]) weekly for eight weeks. Injections (intramuscular) rotated between the quadricep and the tibialis anterior, appropriate for their treatment group. At completion of treatment mice were sacrificed via cervical dislocation and guillotine. Mice were weighed at the beginning and end of treatment. Blood serum was collected and tissues (gastrocnemius, TA, soleus, quadricep) were harvested and flash frozen in liquid nitrogen.

### Serum MMA

MMA was measured on Thermo Q Exactive (Thermo Scientific) as previously described with minor modifications (12). Briefly, 50 µl of serum incubated with internal standard, methyl-d3-malonic acid (D3-MMA), (Toronto Research Chemicals) for 15 min at room temperature. Proteins were then precipitated with 0.4% formic acid in acetonitrile. Supernatant was evaporated in SpeedVac and extracts were derivatized to ester with 3N HCl in n-butanol during incubation at 60°C for 30 min. After drying final extract were reconstituted in 100uL mobile phase for analysis on LCMS system containing Thermo Ultimate 3000 UPLC and Thermo Q Exactive mass spectrometer. Mass spectrometer was operating in positive PRM mode and set to detect m/z 231.16 for MMA and m/z 234.16 for ^13^C_4_-MMA. For quantification m/z 119 was used for MMA and m/z 123 was used for ^13^C_4_-MMA.

### Microbiological *Lactobacillus leichmannii* assay for total vitamin B12

Plasma vitamin B12 concentrations were measured by a modified version of published procedures (13). Briefly, plasma was thawed on ice and diluted 10-fold in extraction buffer containing sodium hydroxide (8.3mM), acetic acid (20.7 mM), and sodium cyanide (0.45mM), pH 4.5 to liberate protein-bound vitamin B12 in plasma. Samples were autoclaved for 10 minutes at 121°C, cooled on ice, and spun down for 15 minutes at maximum speed in a tabletop centrifuge at 4°C. The supernatant was removed, and the extracted volume recorded. A standard curve was prepared using 1 pg/µL, 2 pg/µL, 4 pg/µL, 6 pg/µL, and 8 pg/µL cyanocobalamin standards added to each tube at a final concentration of 0-80 pg. Prior to running each assay, a stock of cyanocobalamin was measured, and the concentration determined using the molar extinction coefficient 27,500 cm^-1^/M at 361nm. Difco B12 Assay Media (BD, #245710) was prepared according to the manufacturer’s instructions. In glass borosilicate tubes, 10mL assay mixtures were prepared for the following: 1) *samples* (5 mL B12 assay media, 40µL extracted plasma, 4.86 mL deionized water), 2) *standards* (5 mL B12 assay media, 50µL diluted cyanocobalamin, 4.85 mL deionized water), 3) *blank* (5 mL B12 assay media, 40µL extraction buffer, 4.86 mL deionized water), and 4) *zero standard* (5 mL B12 assay media, 40µL extraction buffer, 4.86 mL deionized water). Assay mixture tubes were autoclaved for 20 minutes at 121°C and allowed to cool to room temperature before inoculation with 100µL of either 0.9% saline (*blank*) or L. *leichmannii* diluted 1:50 in 0.9% saline (*zero, standards, samples*). Cultures were incubated at 37°C for 24 hours. The following day, tubes were vortexed and subsequently plated in triplicate into 96-well plates (300µL/well), including a standard curve on each plate. *L. leichmannii* growth was quantified at 595 nm by Epoch Microplate Spectrophotometer (Biotek Instruments).

### Uracil in mtDNA Measured by Real-Time PCR

DNA was isolated from frozen mitochondrial pellets using QIAprep Spin Miniprep Kit (Qiagen). Mitochondrial pellets from red muscle and gastrocnemius were collected as previously described (11). Uracil in mtDNA was quantified using a previously described real-time PCR assay (7).

### mtDNA Copy Number and mtDNA Deletion qPCR

DNA from muscle was isolated using High Pure PCR Template Preparation Kit (Roche). DNA concentrations were measured by Nanodrop 2000c Spectrophotometer (Thermo Scientific). Quantitative PCR to measure mtDNA content was performed as previously described (14) values are relative to beta-2-microglobulin (B2M) nuclear levels. Frequency of a mtDNA 3860-base pair deletion was also measured by a previously described quantitative PCR assay (15).

### Uracil in nDNA Measured by gas chromatography-mass spectrometry

DNA was isolated using High Pure PCR Template Preparation Kit (Roche) per the manufacturer’s protocol. Following RNAse A treatment, DNA was purified again using High Pure PCR Purification Kit (Roche) per manufacturer’s protocol, and DNA concentrations were quantified by Qubit (Thermo Scientific). 1.4 µg of DNA was treated with uracil DNA glycosylase (UDG, New England Biolabs, Inc.) for 60 minutes at 37°C with gentle shaking. Samples were derivatized as previously described (16). Uracil levels were quantified by gas chromatography-mass spectrometry (GC-MS) using an Exactive GC mass spectrometer with Trace 1300 GC and TriPlus RSH autosampler (Thermo Electon LLC). In splitless injector mode 3uL of derivatized extract was injected on TG-5SILMS 30m X 0.25mm X 0.25µm column (Thermo). The MS transfer line was set to 280°C and the ion source temperature to 300°C. The MS was operating in negative SIM mode with chemical ionization using methane at flow 1.9 mL/min. AGC target was set to 5e5, max IT 200ms, resolution 30,000, emission current 50µA, electron energy 70eV. Uracil was quantified using m/z 337.04 in linear calibration range 1pg to 32 pg.

### Mitochondrial Mass

Mitochondrial mass in tibialis anterior and soleus was measured according to the manufacturer’s instructions using the Citrate Synthase Activity Assay Kit (Sigma) and normalized to protein concentration. Protein concentrations were assessed by Lowry–Bensadoun assay (17). Absorbance of the colorimetric assay was quantified by Epoch Microplate Spectrophotometer (Biotek Instruments).

### Respirometry in Frozen Samples

The oxygen consumption rate (OCR) of tibialis anterior was measured using the Seahorse XFe24 and red muscle was measured using the Seahorse XFe96 Extracellular Flux Analyzer (Agilent Technologies) as previously described with minor modifications (18). Briefly, mitochondria from frozen muscle were isolated in ice-cold MAS buffer using a Potter–Elvehjem (Teflon-glass) homogenizer with 25 strokes followed by centrifugation. Protein concentrations of the mitochondrial supernatant were determined by a Pierce BCA Protein Assay (Thermo Fisher). Mitochondrial homogenates were loaded into the assay plates and centrifuged at 2,000×g for 5 min at 4°C with no brake. For TA in the Seahorse XFe24 microplate we loaded 150 µg of homogenate in 100 µl of MAS. For red muscle using the Seahorse XFe96 we loaded 30 µg of homogenate in 70 µl of MAS. The OCR of each complex was determined using Wave software (Agilent) and was defined by the highest respiratory capacity value following the injection of the corresponding complex stimulating substrate: complex I, 1 mM NADH; complex II, 5 mM succinate + 2 µM rotenone; and complex IV, 0.5 mM N,N,N′,N′-Tetramethyl-p-phenylenediamine dihydrochloride (TMPD) + 1 mM ascorbic acid. For baseline, mix time was 2 min and measure 3 min. For substrates, mix and measure times were 1 and 3.5 min, respectively. A 2-min wait time was included for oligomycin-resistant respiration measurements.

### Immunoblotting

Tissues were lysed by sonication in lysis buffer (150 mM NaCl, 10 mM Tris–Cl, 5 mM EDTA pH 8, 5 mM dithiothreitol, 1% Triton X-100, protease inhibitor), and cell debris was removed by centrifugation at 4°C for 10 min at 14,000×g. Protein concentrations of tissue lysate supernatants were assessed by Lowry-Bensadoun assay. Samples were boiled with SDS–PAGE sample loading buffer (6× SDS), and 30 μg of protein was loaded to each well of a 10% Tris– glycine SDS–PAGE gel (BioRad). Proteins were transferred to a PVDF membrane (Millipore). Membranes were blocked with 5% (wt/vol) nonfat dry milk in 1× PBS with 0.05% Tween-20 for 1 h. Membranes were then incubated for 1 h with 1:2,000 primary antibody [glyceraldehyde-3-phosphate dehydrogenase (GAPDH), Cell Signaling #14C10; α-MTR, ProteinTech #25896; COX IV (3E11), Cell Signaling #4850] in 5% bovine serum albumin (BSA) 0.02% NaN3. Membranes were then washed three times in 1× PBS with 0.01% Tween-20 and incubated for 1 h with HRP-conjugated secondary antibody diluted 1:20,000 in 5% (wt/vol) nonfat dry milk in 1× PBS with 0.05% Tween-20. Membranes were exposed with chemiluminescent substrate (BioRad) and visualized using a ProteinSimple Imager (Bio-techne). Protein levels were quantified using ImageJ software.

## Results

### B12-deficient diet lowers plasma B12 and elevates MMA in the *Mtr^+/−^* mouse model

There were no differences in mouse body weight between diet groups or as a result of *Mtr* genotype at any timepoint (Figure S1A). Plasma vitamin B12 concentration, as measured by *Lactobacillus leichmannii* microbiological assay, was reduced to 1/20^th^ of control levels (*p*=0.0001) in response to -B12 diet (Figure S1B). Serum methylmalonic acid (MMA) concentrations, as measured with LC-MS/MS, were elevated by 20% (*p*=0.0006) with -B12 diet exposure (Figure S1C). These biomarker changes demonstrate a reduction in B12 status in mice consuming the -B12 diet. Soleus muscle weight was reduced (*p=*0.04) in control mice consuming -B12 diet (Figure S1G). Interestingly, gastrocnemius muscle weight was significantly reduced (*p=*0.03) with reduced *Mtr* expression (Figure S1F).

### Reduced *Mtr* expression and dietary B12 deficiency alter oxidative capacity of skeletal muscle

We examined TA and red muscle as two distinct skeletal muscle groups in which to compare the effects of reduced *Mtr* expression and B12 deficiency on mitochondrial outcomes. Generally, the TA is comprised of mainly glycolytic myofibers (type IIB) while red muscle is comprised of predominantly oxidative myofibers (type I, IIA) (19). To explore the effects of reduced *Mtr* expression and B12 deficiency on mitochondrial oxidative capacity, in type IIB muscle like TA, maximal respiratory complex activity was quantified. Reduced *Mtr* expression significantly decreased maximal activity of complex I (*p*=0.01), complex II (*p*=0.001), and complex IV (*p*=0.006) (**Figure 2**A-C) in TA. Further, B12-deficient diet coupled with reduced *Mtr* expression decreased maximal activity of complex IV (*p*=0.04, Figure 2C), suggesting that *Mtr* genotype and B12 deficiency impair oxidative capacity of TA muscle mitochondria. MTR protein levels in TA were reduced by 80% (*p*<0.0001) in *Mtr^+/−^* mice on the C diet (Figure 2D-E). Interestingly, TA MTR protein levels were also reduced (*p*=0.0001, for effect of diet) as a result of -B12 diet exposure (Figure 2D-E). Cytochrome c oxidase subunit 4 isoform 1 (COX IV) protein levels in TA decreased (*p*=0.005) in response to B12-deficient diet (Figure 2D, F), paralleling the effect of the -B12 diet on complex IV activity (Figure 2C).

**Figure 2.**
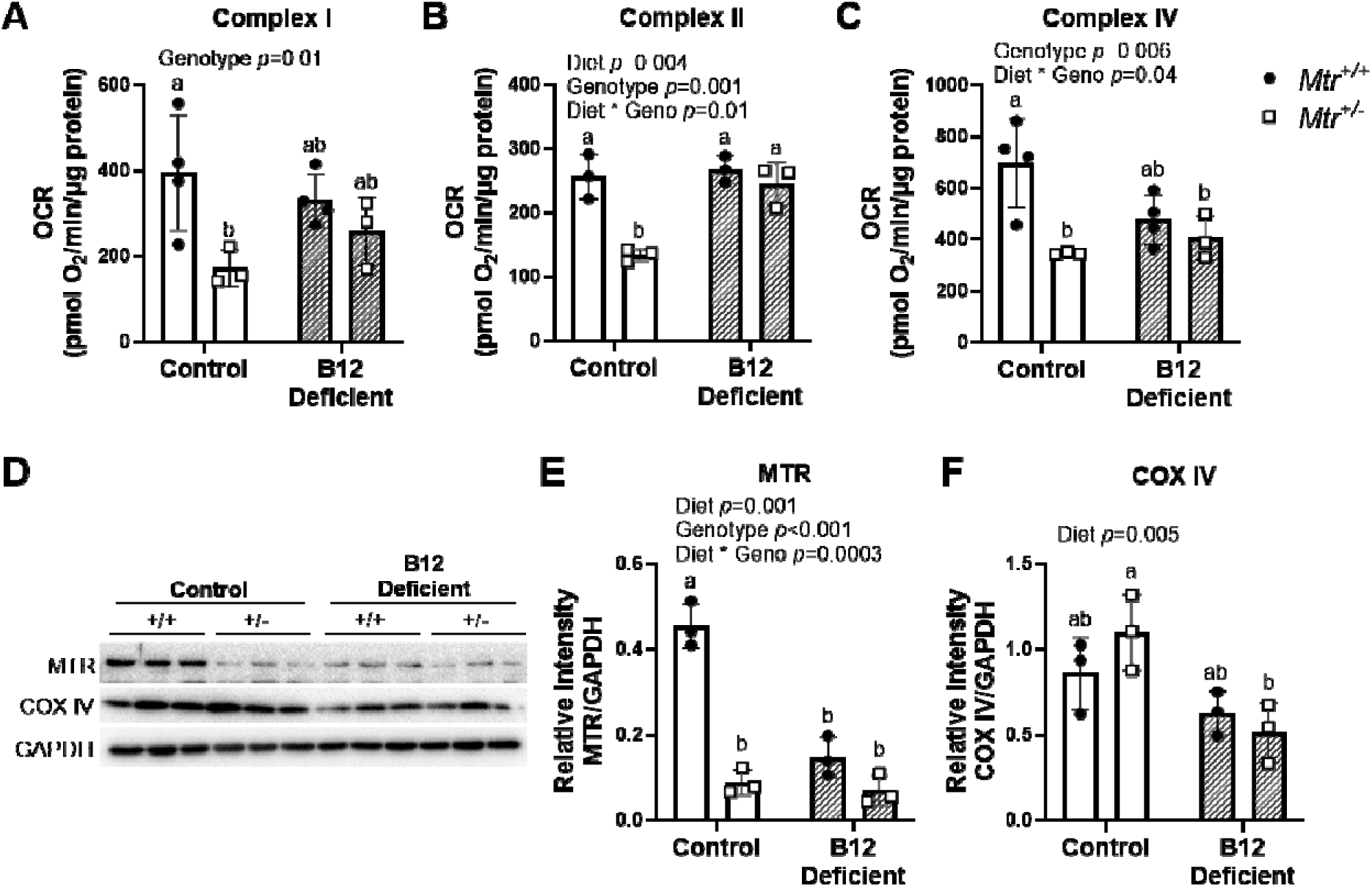
Impaired functional capacity of mitochondrial electron transport chain complexes in tibialis anterior muscle. A) Complex I activity, B) complex II activity, and C) complex IV activity using respirometry in tibialis anterior homogenates; n=3-4 per group. (D) Western blot of TA MTR and COX IV expression; n=3 per group. (E) TA MTR protein levels normalized to GAPDH; n=3 per group. (F) TA COX IV protein expression normalized to GAPDH; n=3 per group. Data are shown as mean ± SD and were analyzed by two-way ANOVA with Tukey’s post hoc analysis, with significance defined as *p* ≤ 0.05. Residual analyses were performed to check the model assumptions of normality and homogeneous variance. Groups not connected by a common letter are significantly different. COX IV, complex IV; GAPDH, glyceraldehyde-3-phosphate dehydrogenase; MTR, methionine synthase; OCR, oxygen consumption rate.

In red muscle, maximal respiratory complex activity of complexes I and IV was also measured. Due to tissue input limitations, complex II was not measured. B12-deficient diet modestly reduced activity of complex I (*p*=0.02) and complex IV (*p*=0.03) in red muscle (**Figure 3**A-B). This suggests dietary B12 deficiency may have a stronger effect than reduced *Mtr* expression on reducing mitochondrial respiratory capacity in more oxidative tissue like red muscle, compared to glycolytic muscle such as TA.

**Figure 3.**
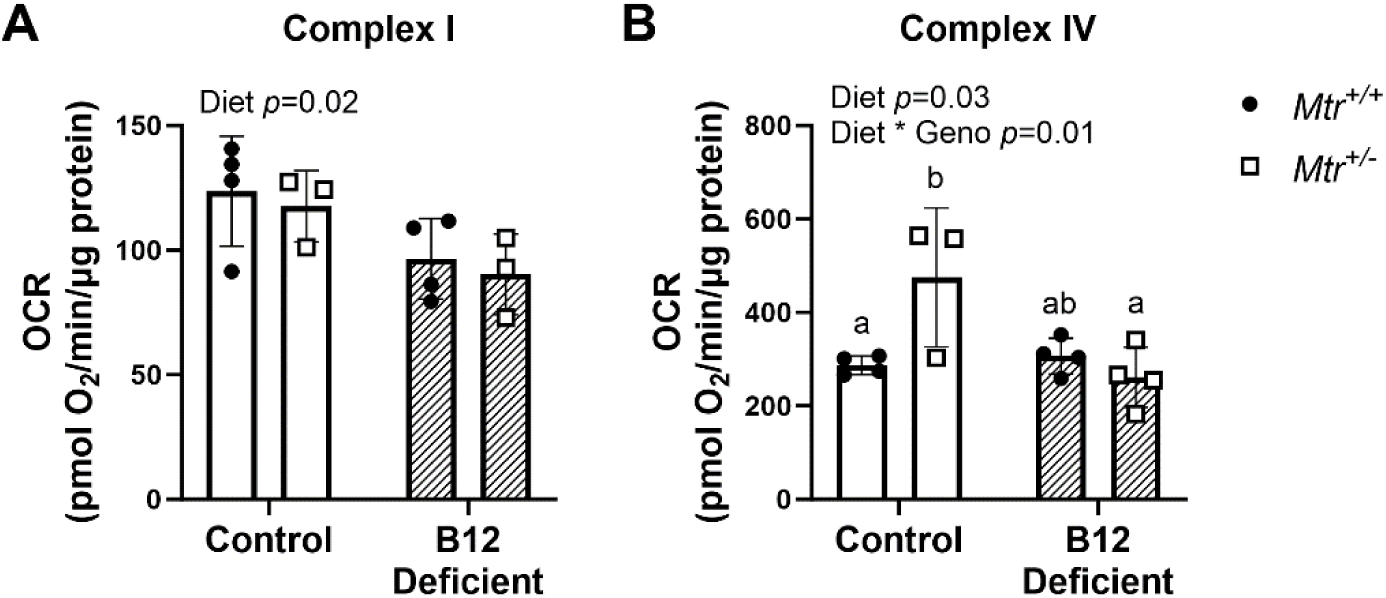
Impaired functional capacity of mitochondrial electron transport chain complexes in red muscle. A) Complex I activity and B) complex IV activity using respirometry in red muscle homogenates; n=3-4 per group. Data are shown as mean ± SD and were analyzed by two-way ANOVA with Tukey’s post hoc analysis, with significance defined as *p* ≤ 0.05. Residual analyses were performed to check the model assumptions of normality and homogeneous variance. Groups not connected by a common letter are significantly different. MTR, methionine synthase; OCR, oxygen consumption rate.

### Dietary B12 deficiency leads to uracil misincorporation mtDNA, but does not impact mtDNA content, mitochondrial mass, or mtDNA deletion frequency in skeletal muscle

Because of the role of B12 in FOCM to support nucleotide synthesis, we wanted to examine whether disruptions to cytosolic dTMP synthesis by dietary B12 deficiency or by *Mtr^+/−^* genotype would affect mitochondrial DNA (mtDNA) integrity. We previously developed an assay to quantify uracil misincorporation into mtDNA (7). As described earlier, uracil misincorporation arises when *de novo* dTMP synthesis is impaired by either folate or B12 deficiency (5,20). To assess whether reduced *Mtr* expression or dietary B12 deficiency affected mtDNA integrity, uracil content of red muscle mtDNA was measured by quantitative real-time PCR assay (7). Uracil levels in mtDNA were elevated by >10-fold in mice consuming the -B12 diet compared to mice consuming the C diet (*p*=0.04) (**Figure 4**). *Mtr* genotype did not significantly alter mtDNA uracil levels in red muscle or gastrocnemius mitochondria (Figure 4A-B). Elevated uracil accumulation (*p*<0.001) in mtDNA was also observed with -B12 diet in whole gastrocnemius muscle mitochondria (Figure 4B). In addition, the phenotype of B12 deficiency leading to uracil in skeletal muscle was unique to mtDNA, as no change was observed in quadricep nDNA (**Supplementary Figure 2**) as a result of either *Mtr^+/−^* genotype or dietary B12 deficiency in this model.

**Figure 4.**
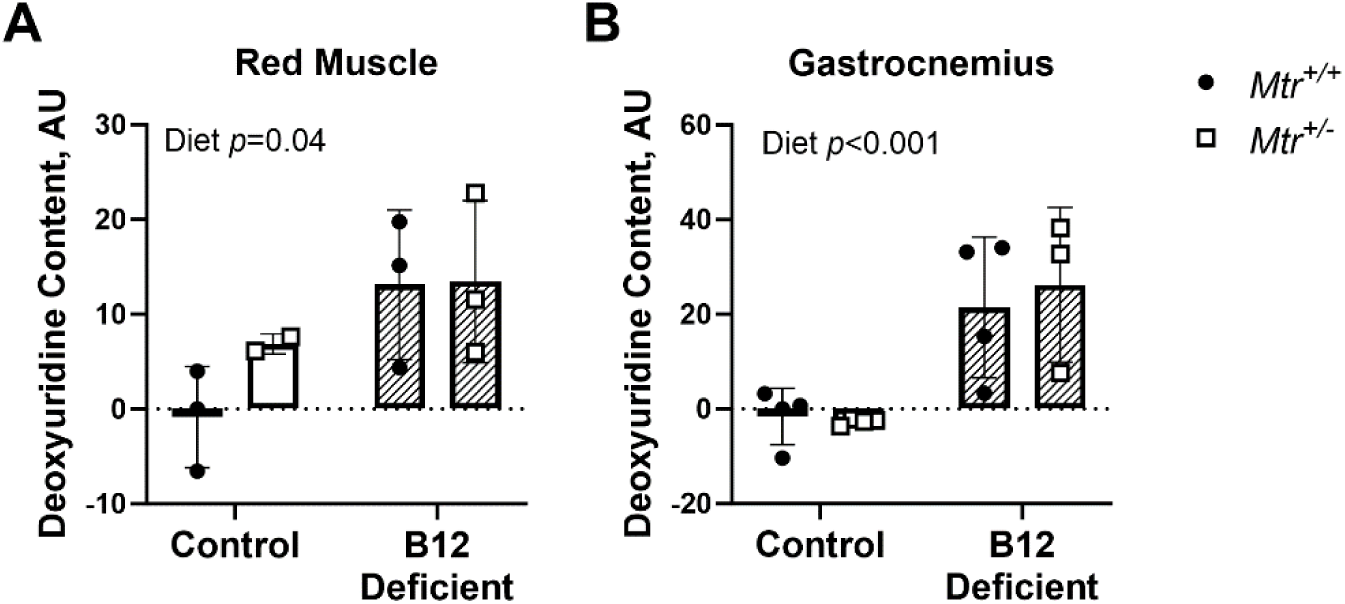
B12 deficiency increases uracil in skeletal muscle mtDNA. **(A)** Uracil content in red muscle mitochondrial DNA; n=2-3 per group. **(B)** Uracil content in gastrocnemius muscle mitochondrial DNA; n=2-3 per group. Data are shown as mean ± SD and were analyzed by two-way ANOVA with Tukey’s post hoc analysis, with significance defined as *p* ≤ 0.05. Residual analyses were performed to check the model assumptions of normality and homogeneous variance. AU, arbitrary units; mtDNA, mitochondrial DNA; *Mtr*, methionine synthase.

To understand how uracil in mtDNA may relate to mitochondrial function, we investigated multiple mitochondrial markers. mtDNA content, the abundance of mtDNA relative to nuclear DNA (nDNA), is increasingly recognized as an indicator of mitochondrial function (9,21). mtDNA content in TA and red muscle were unaffected by *Mtr^+/−^* genotype or exposure to the -B12 diet (**Figure 5**A-B), consistent with previous findings (7). Mitochondrial mass was also unchanged, as there was no difference in citrate synthase activity by exposure to the -B12 diet or *Mtr^+/−^* genotype (Figure 5C-D). There is a 3,680 base pair deletion in murine mtDNA that is analogous to the common 4,977 base pair deletion in human mtDNA (15). The frequency of these mutations is known to increase with elevations in oxidative stress (reactive oxygen species) and with age (15). However, neither *Mtr^+/−^* genotype nor B12-deficient diet altered the frequency of the mtDNA deletion (Figure 5E-F). This suggests that uracil in mtDNA does not affect mitochondrial function by altering mtDNA content, mitochondrial mass, or mtDNA mutation frequency.

**Figure 5.**
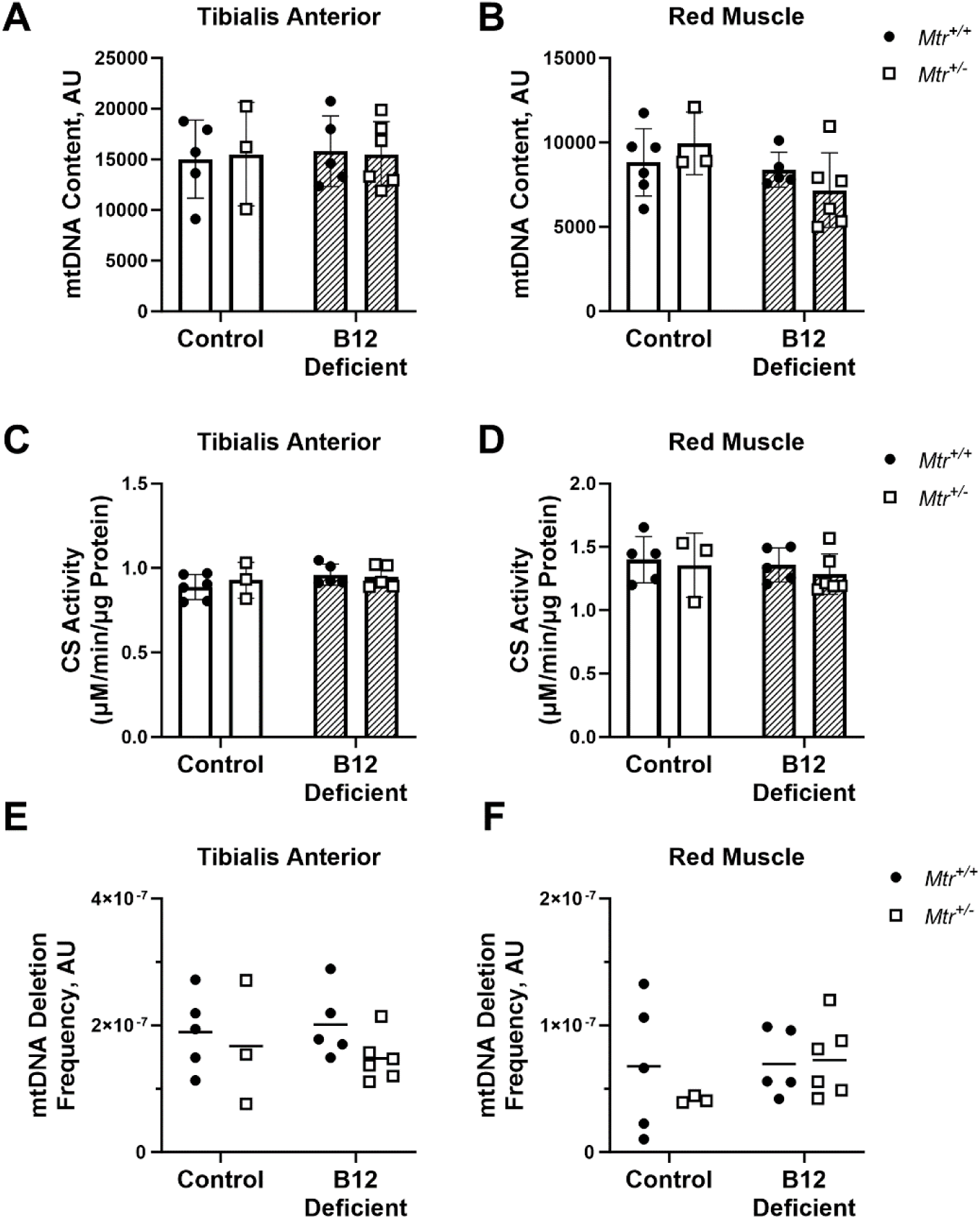
Biomarkers of mitochondrial DNA content and mass in skeletal muscle. A) Tibialis anterior and B) red muscle mtDNA copy number determined using real-time PCR; n = 3 6 per group. C) Tibialis anterior and D) red muscle mitochondrial mass quantified using citrate synthase activity; n = 3-6 per group. E) Tibialis anterior and F) red muscle mtDNA common deletion frequency determined using real-time PCR; n=3-6 per group. Data are shown as mean ± SD and were analyzed by two-way ANOVA with Tukey’s post hoc analysis, with significance defined as *p* ≤ 0.05. Residual analyses were performed to check the model assumptions of normality and homogeneous variance. AU, arbitrary units; CS, citrate synthase; mtDNA, mitochondrial DNA; *Mtr*, methionine synthase.

### B12 supplementation improves complex IV maximal capacity in aged mice

In older adults, a common physiological change of aging is the loss of skeletal muscle mass, strength, and function. Additionally, age-related skeletal muscle decline is closely linked to mitochondrial dysfunction (22), a key hallmark of aging (23,24). Therefore, to explore the potential therapeutic benefits of B12 supplementation on mitochondrial function we measured maximal oxidative capacity of gastrocnemius in aged mice. A 2-fold increase in maximal activity of complex IV was observed in aged mice receiving the B12 supplement compared to non-supplemented controls (*p*=0.04) (**Figure 6**B).

**Figure 6.**
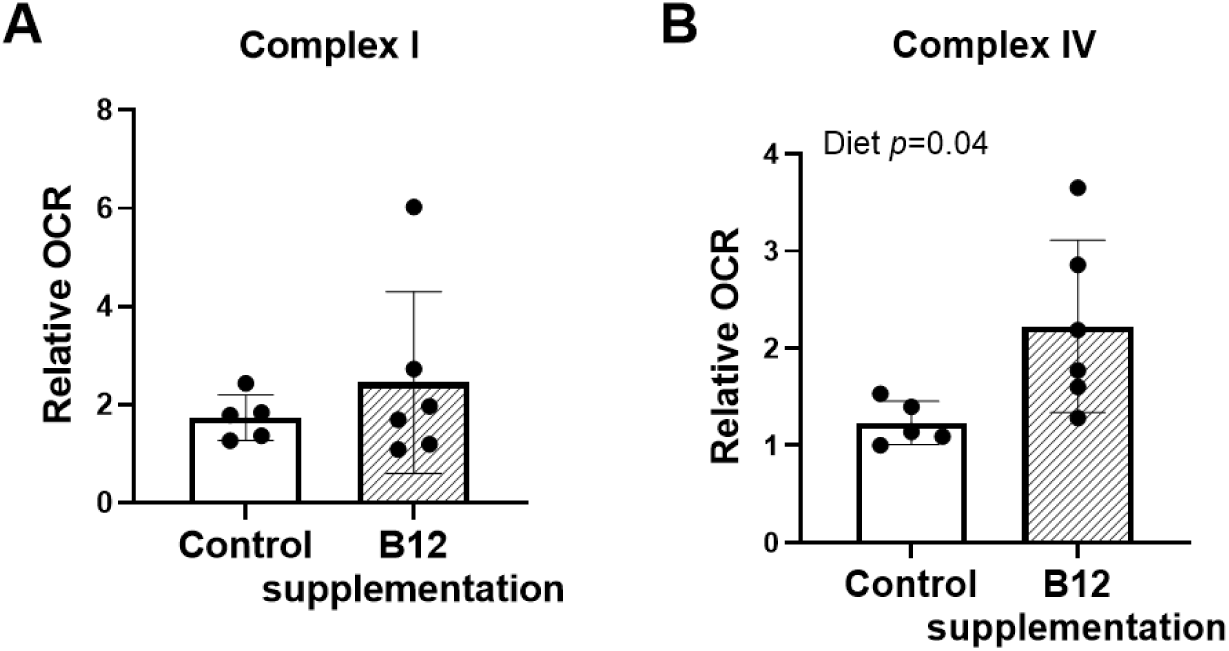
Improved functional capacity of mitochondrial electron transport chain complexes in gastrocnemius muscle of aged mice given B12 supplementation for 8 weeks. A) complex I activity and B) complex IV activity using respirometry in gastrocnemius homogenates; n=4-5 per group. Samples were run across several plates, and data were normalized to the lowest value on each plate. Data are shown as mean ± standard deviation and were analyzed using a student’s t-test, significance defined as *p* ≤ 0.05. Control, control/saline treatment; OCR, oxygen consumption rate; Supplementation, treated with 1.25 µg B12 per week.

## Discussion

In this study, the *Mtr^+/−^* mouse model was used to determine how impaired FOCM (and the addition of dietary B12 deficiency) affect mitochondrial function and mtDNA integrity. Additionally, we investigated whether B12 supplementation could mitigate age-related decline in mitochondrial function. As mentioned earlier, B12 is required by only two enzymes: MTR and MCM. Elevated MMA, a by-product of excess MCM substrate, is a classical marker to diagnose B12 deficiency. It has been previously shown that the *Mtr^+/−^* mouse model does not exhibit altered plasma MMA levels (8), meaning that the model recapitulates one effect of B12 deficiency that is specific to impaired FOCM. Reduced *Mtr* expression led to impaired oxidative capacity and accumulation of uracil in mtDNA in mouse liver (8). Here, control and B12-deficient diets were used in this study to compare the effects of systemic B12 deficiency to the FOCM-specific effects of B12 deficiency modeled by *Mtr^+/−^*.

In TA, MTR protein levels were decreased with *Mtr^+/−^*genotype and dietary B12 deficiency, confirming lower *Mtr* expression in skeletal muscle (Figure 2D-E). *Mtr^+/−^* genotype led to lower respiratory capacity in three electron transport chain complexes in TA (Figure 2A-C). This agrees with previous findings where *Mtr^+/−^* genotype impaired respiratory capacity in mouse liver tissue (8). Mice consuming the B12-deficient diet also exhibited lower respiratory capacity in TA, but only in complex IV (Figure 2C). Interestingly, red muscle respiratory capacity was unaffected by *Mtr^+/−^*genotype, showing only a modest effect from the -B12 diet (Figure 3A). This suggests some muscle groups may be more sensitive to reduced *Mtr* expression or dietary B12-deficiency.

The elevation of uracil in mtDNA in red muscle and gastrocnemius (Figure 4) in mice consuming the B12-deficient diet indicates that in skeletal muscle, dietary B12-deficiency impairs cytosolic dTMP synthesis and this in turn hinders mtDNA replication. The increase in mtDNA uracil as a result of exposure to the B12-deficient diet was not accompanied by an increase in uracil in nDNA (Supplementary Figure 2), suggesting that mtDNA is more sensitive to the effects of B12 deficiency than is nDNA, corroborating our previous finding in liver tissue (7,8). It is worth noting that the dietary B12 deficiency in this model is relatively mild, given the modest increase in serum MMA (Supplementary Figure 1C). Although the effect of *Mtr^+/−^*genotype on elevating uracil in mtDNA in skeletal muscle did not reach statistical significance, it is not clear whether this is due to limited sample size or true biological effect. This relatively limited sample size is one limitation of the study.

Mitochondrial dysfunction is a recognized hallmark of the aging process (23,24). Aging is associated with a decline in mitochondrial biogenesis and impaired OXPHOS activity (25–27). In skeletal muscle, these mitochondrial impairments manifest as decreased ATP availability, compromised muscle fiber maintenance, and increased oxidative damage (28). B12 deficiency has similarly been linked to muscle weakness, mitochondrial dysfunction, and increased susceptibility to frailty-related outcomes in older adults (22,29). In this study we investigated the effects of an eight-week regimen of intramuscular B12 injections in aged male mice. The results showed a two-fold increase in mitochondrial complex IV activity in gastrocnemius muscle as a result of B12 treatment (Figure 6B). Although skeletal muscle loss associated with mitochondrial dysfunction is frequently observed in older adults (22), the underlying mechanisms remain poorly understood. One plausible explanation involves the critical role of B12 in supporting dTMP synthesis and maintening DNA integrity. Given that older adults are at increased risk of low B12 status due to decreased absorption (1), our findings suggest that B12 supplementation may be a promising strategy to improve mitochondrial function and restore maximal oxidative capacity in aging skeletal muscle.

This study is the first to demonstrate that B12 deficiency impairs maximal respiratory capacity and leads to uracil misincorporation into skeletal muscle mtDNA. Additionally, we provide the first evidence that B12 supplementation enhances mitochondrial oxidative phosphorylation complex activity. *De novo* dTMP synthesis is compartmentalized, that means it happens distinctly in the cytosol, mitochondria, and nuclear compartments. Previous work has shown that nuclear *de novo* dTMP synthesis is relatively protected from impairements in FOCM through multiple mechansims (5), leaving the mitochondrial dTMP pool more vulnerable to FOCM impairment. Collectively, our data demonstrates that low B12 status reduces cytosolic dTMP availability, resulting in uracil misincorporation into mtDNA and impaired OXPHOS complex activity; conversely, B12 supplementation restores complex activity in aging skeletal muscle. These findings are also consistent with observations that disruptions in cytosolic dTMP synthesis lead to mtDNA instability and impaired mitochondrial function (7,8). Further, mtDNA content, mitochondrial mass, and mtDNA deletion frequency were not affected by *Mtr^+/−^* genotype or B12 deficiency, suggesting that these markers appear to be independent of mtDNA uracil accumulation or complex respiratory capacity in skeletal muscle (Figure 5). Although dTMP pools in postmitotic tissues are generally thought to be maintained primarily through the dTMP salvage pathway synthesis (30), the presence of uracil in mtDNA suggests a substantial contribution from cytosolic *de novo* dTMP synthesis to total mitochondrial dTMP availability in skeletal muscle. Supporting this, previous studies in cell culture models have shown that mitochondrial dTMP synthesis alone is insufficient to maintain adequate mitochondrial dTMP pools and prevent uracil misincorporation into mtDNA (31). Further research is needed to clarify the underlying mechanisms and to explore the potential of mtDNA uracil as a biomarker for mitochondrial dysfunction and age-associated skeletal muscle decline.

## Supporting information

supplementary revised fig 2

## Acknowledgments

The authors’ responsibilities were as follows. KEH, LFC, AET, MSF: designed the research; KEH, LFC, ARW, WM, NMV, OVM: performed research and analyzed data; KEH, LFC, ARW, WM, OVM, NMV, AET, MSF: wrote the paper; MSF: had primary responsibility for the final content; all authors have read and approved the final manuscript. The authors acknowledge assistance with statistical analysis from the Cornell Statistical Consulting Unit and assistance from Ivan Keresztes with use of the Exactive GC mass spectrometer.

## Supplementary Figures

**Supplementary Figure 1.**
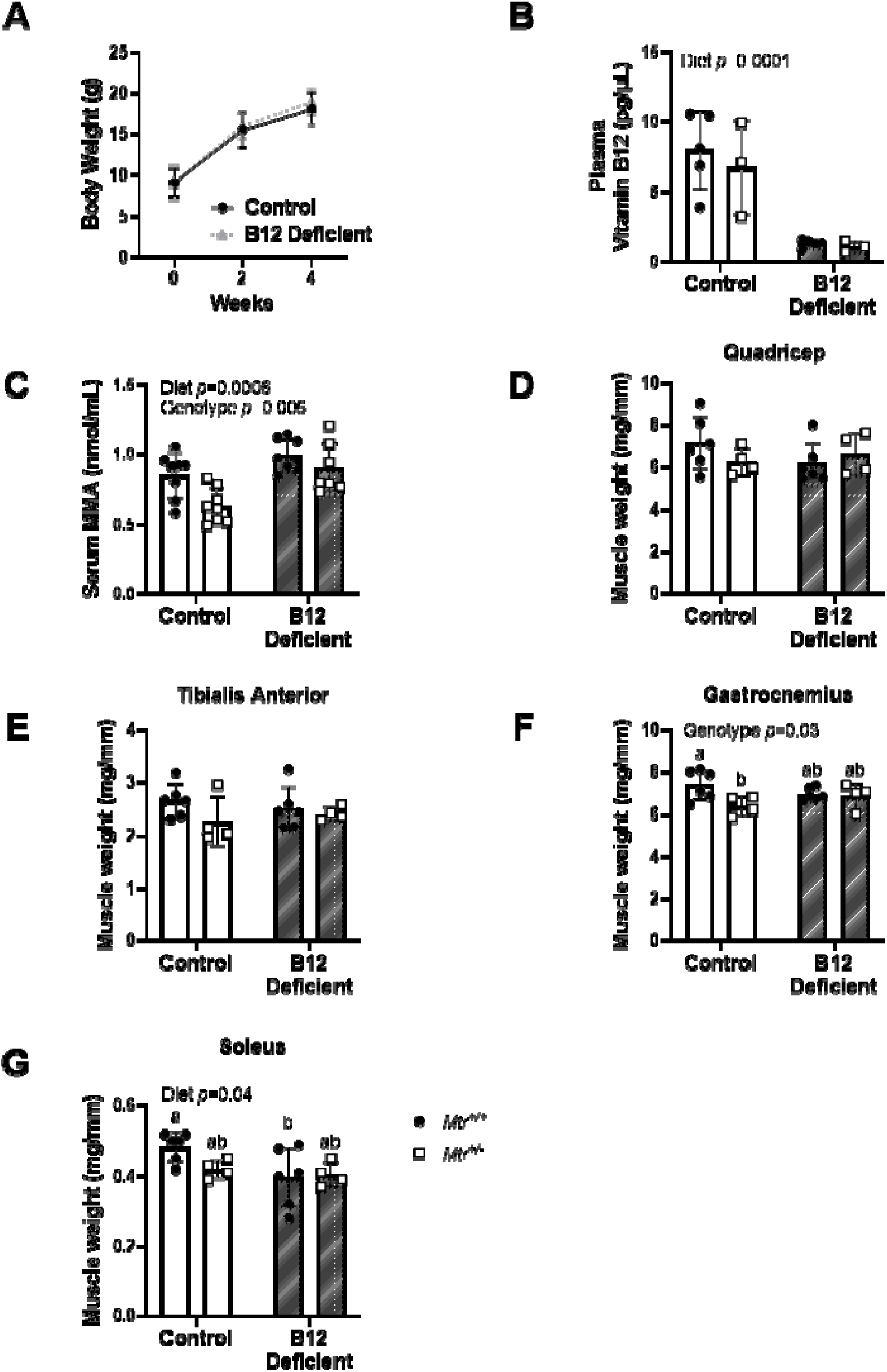
Body weight, plasma B12 measurement, and muscle weights. (A) At weaning, mice were placed on the control or B12 deficient diet and body weights were measured at weaning and every two weeks thereafter; n = 17-18 per group. Two-way mixed ANOVA was used to assess fixed effects of time, diet, and time-diet interaction. (B) Plasma vitamin B12 concentrations measured by *Lactobacillus leichmannii* microbiological assay; n= 3-5 per group. (C) Serum methylmalonic acid concentrations measured by LC-MS/MS; n=7-8 per group. Residual analyses were performed to check the model assumptions of normality and homogeneous variance. Data are shown as mean ± s.d. and significance was defined as *p* ≤ 0.05. MMA, methylmalonic acid. (D-G) Muscle weights (mg) from quadricep, tibialis anterior, gastrocnemius, and red muscle were normalized to tibia length (mm); n=4-6 per group. Two-way ANOVA with Tukey’s post-hoc analysis, with significance defined as *p* ≤ 0.05. Groups not connected by a common letter are significantly different. MMA, methylmalonic acid; *Mtr*, methionine synthase.

**Supplementary Figure 2.**
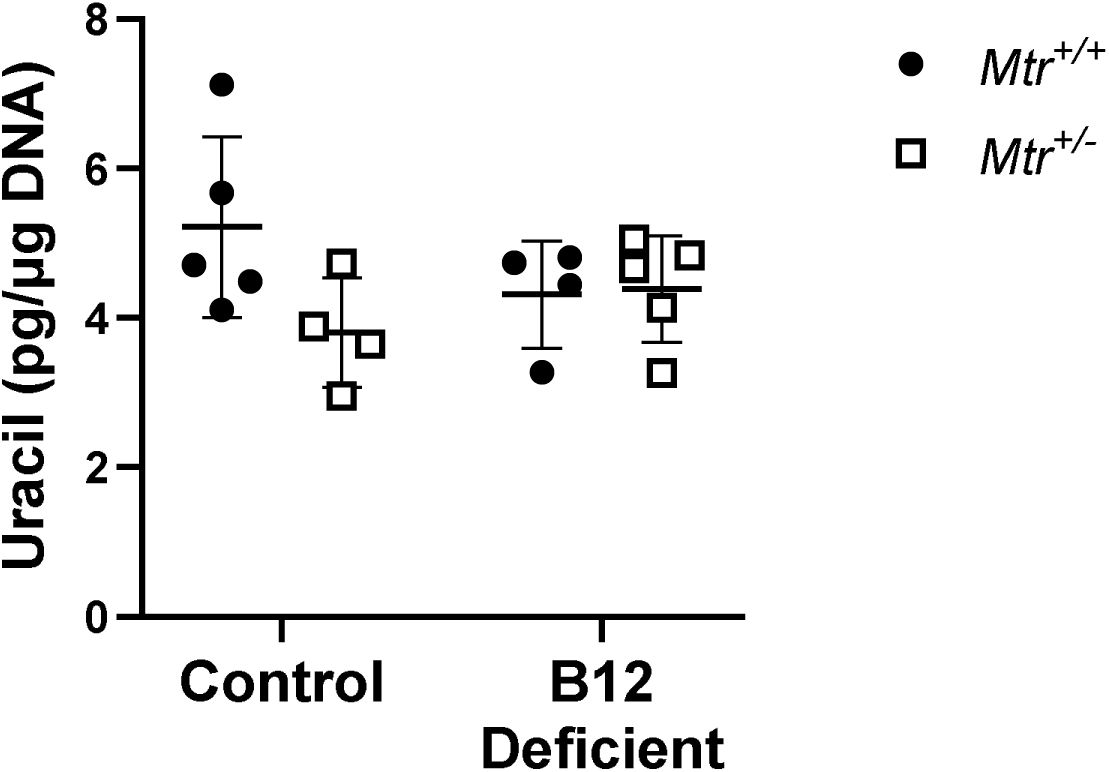
Genomic Uracil in Quadricep. Genomic uracil measured by GC/MS n=4-5 per group. Two-way ANOVA with Tukey’s post hoc analysis, with significance defined as *p* ≤ 0.05.

