## supplementary revised fig 2 for "Vitamin B12 supports skeletal muscle oxidative phosphorylation capacity in male mice"

**Supplementary Figures:**

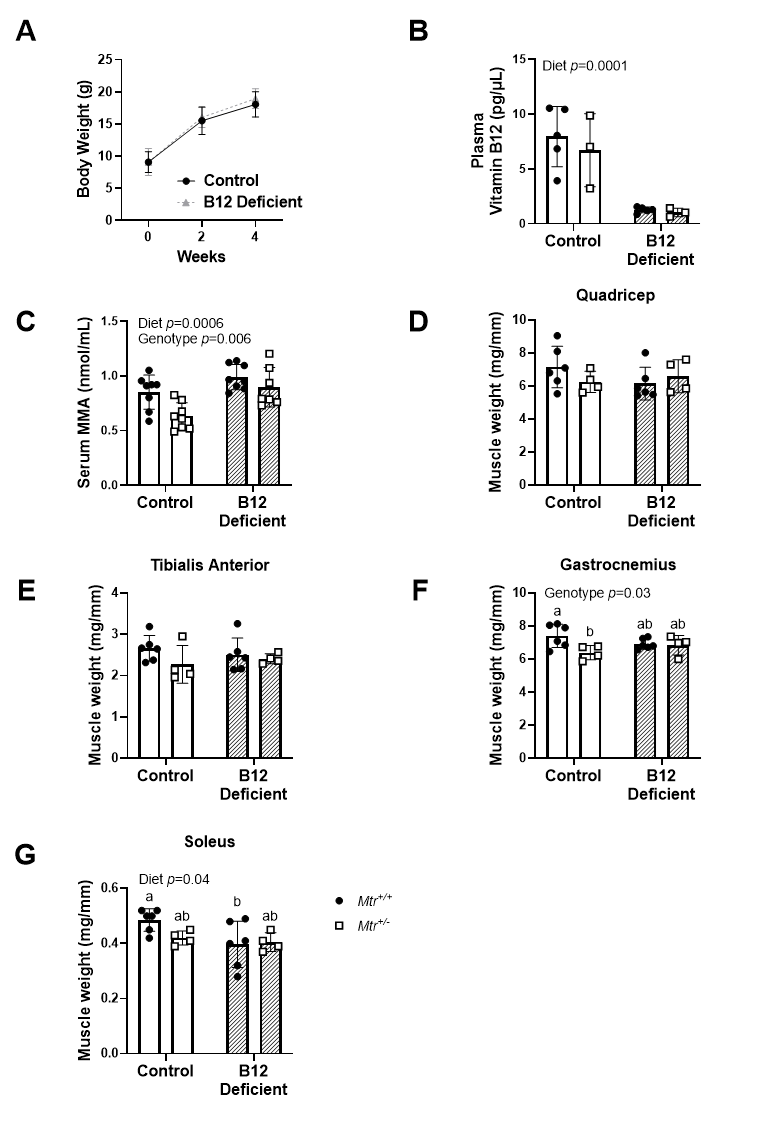

**Supplementary Figure 1. Body weight, plasma B12 measurement, and muscle weights.** (A) At weaning, mice were placed on the control or B12 deficient diet and body weights were measured at weaning and every two weeks thereafter; n = 17-18 per group. Two-way mixed ANOVA was used to assess fixed effects of time, diet, and time-diet interaction. (B) Plasma vitamin B12 concentrations measured by *Lactobacillus leichmannii* microbiological assay; n= 3-5 per group. (C) Serum methylmalonic acid concentrations measured by LC-MS/MS; n=7-8 per group. Residual analyses were performed to check the model assumptions of normality and homogeneous variance. Data are shown as mean ± s.d. and significance was defined as *p* ≤ 0.05. MMA, methylmalonic acid. (D-G) Muscle weights (mg) from quadricep, tibialis anterior, gastrocnemius, and red muscle were normalized to tibia length (mm); n=4-6 per group. Two-way ANOVA with Tukey’s post-hoc analysis, with significance defined as *p* ≤ 0.05. Groups not connected by a common letter are significantly different. MMA, methylmalonic acid; *Mtr*, methionine synthase.

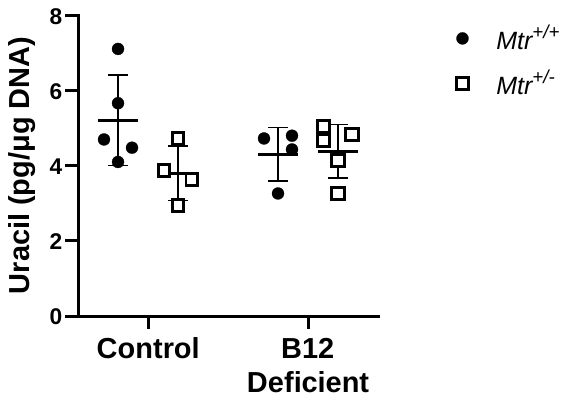

**Supplementary Figure 2. Genomic Uracil in Quadricep.** Genomic uracil measured by GC/MS n=4-5 per group. Two-way ANOVA with Tukey’s post hoc analysis, with significance defined as *p* ≤ 0.05.
